# Targeting skin ageing at the source: a novel cellular reprogramming approach using natural extract combinations

**DOI:** 10.64898/2026.10.13.758751

**Authors:** Maria L. Perciato, Merlin Davies, Pouria Akhbari, Hubert Denise, Adam Clauss, Sarah L. Cole, Kaitlin M. Haas, Brittany Lee, Chantal Jouy, Carine Ballihaut, Ben P. Lee, Lorna W. Harries

**Author notes:** Corresponding author: Professor Lorna W. Harries University of Exeter Medical School Barrack Road, Exeter, EX2 5DW, 44 1392 406773. Equal contributions (Maria L. Perciato, Merlin Davies, Pouria Akhbari).

## Abstract

Cellular senescence is a driver of skin ageing that leads to structural deterioration and aesthetic changes. This study evaluated a novel senomorphic strategy, using synergistic combinations of natural small molecules to target RNA splicing regulation via upstream ERK/AKT signaling pathways.

Nature-identical synthetic molecules, and corresponding extracts were assessed for cytotoxicity, impact on senescent cell load, proliferation, DNA damage load and SASP markers in naturally aged human primary skin cells. A topical, simple formulation was evaluated for impact on senescence and skin structure using 3D human ex vivo full-thickness skin explants exposed to 1.5 J/cm^2^ UVC irradiation.

Combination of *H. lupulus* and *G. biloba* extracts produced a dose-dependent senomorphic impact of up to 45% in aged dermal fibroblasts and 10% in aged human primary keratinocytes (p ≤ 0.0001 & 0.001), without inducing cell proliferation. A 50% decrease in DNA damage load was also observed (p ≤ 0.0001). Qualitatively, the simple formulation mitigated UV-induced epidermal hyperplasia, restored collagen fiber organization, and reduced markers of senescence and inflammation.

Targeting cellular senescence is an emerging traction point to address the root biological causes of skin ageing. Targeting the splicing dysregulation/senescence axis using combinations of natural plant extracts may provide a novel mode of action to reprogram senescent cells for repair and protection of skin structure and function

## Introduction

Skin quality reflects systemic ageing and healthy longevity. Skin ageing is driven by intrinsic and extrinsic factors, such as oxidative stress, UV damage and exposure to pollutants, all of which are potent inducers of cellular senescence (Thau et al., 2025). Senescence, a stable cell-cycle arrest characterized by an altered secretome (SASP, senescence-associated secretory phenotype), plays important physiological roles including wound healing, cancer prevention and embryonic development (Demaria et al., 2014). Failure of immune responses in ageing tissues results in accumulation of senescent cells, which in turn exacerbates chronic inflammation leading to profound changes to the tissues and organs (Campisi and d’Adda di Fagagna, 2007; Wang et al., 2009). Senescence in skin drives changes to cellular, molecular and structural components of the dermis and epidermis (Campisi, 1998), while stem cell exhaustion lowers the regenerative capacity of skin tissue as aged stem cells exhibit compromised ability to repopulate and differentiate following damage or senescence (Demaria et al., 2015). The secretion of the SASP drives aberrant tissue remodeling and extracellular matrix dysfunction, causing alterations in collagen composition and structure, destruction of elastin as well as inflammatory infiltration and induction of fibrosis (Zheng et al., 2026). Collectively, these phenomena lead to the characteristic aesthetic changes associated with ageing, including wrinkles, epidermal hyperplasia, pigmentation changes, skin thinning and deterioration of the underlying skin substructure. Even a small number of senescent cells can be the origin of these damages because of the long range, pleiotropic impact of paracrine (bystander) senescence (Campisi, 1998). Studies have demonstrated that reduction in senescent cells of only 30-35% is enough for clinical benefit (Tchkonia et al., 2013;

Zhu et al., 2021). Evidence is now unequivocal that removal of senescent cells yields clinical benefit in multiple organ systems in animals (Childs et al., 2017; Kiourtis et al., 2024) and in humans (Justice et al., 2019). Senescent cells therefore represent a tractable and emerging target for new therapies aiming to modulate ageing phenotypes.

Senescent cells can be targeted by two approaches; senolytic approaches whereby senescent cells are killed selectively, or senomorphic or reprogramming approaches whereby their deleterious aspects are attenuated (Zhang et al., 2023). Both are valid approaches, and one may have advantages over the other depending on context. Senescence is not a single entity; there are multiple types of senescent cells (Jang et al., 2026), some of which are beneficial (Tripathi et al., 2021). First generation senolytic molecules may not differentiate between these cell types and wholesale ablation of senescent cells could compromise tissue remodeling (Born et al., 2023). The use of senomorphic approaches avoids these disadvantages and allows retention of the cells in their correct tissue context. Several approaches to achieve senomorphic change have been described, including the use of the mTOR inhibitor rapamycin, the AMPK activator metformin or the JAK/STAT inhibitor Ruxolitinib (Saliev and Singh, 2025). These approaches are promising but require long term or intermittent dosing as cessation of treatment may result in rebound activity of SASP factors (Saliev and Singh, 2025). Most of these approaches provide molecular damage control, acting primarily to mute the inflammatory outputs of senescent cells. Reprogramming approaches conversely act upstream in the molecular cascade, and act to bring about a degree of cellular rejuvenation, DNA damage repair and reduction in biological age. Because these approaches correct, rather than mute underlying senescence, the effect of reprogramming often persists beyond cessation of treatment (Latorre et al., 2017). We previously identified restoration of aberrant RNA processing as a novel point of cellular reprogramming (Donega et al., 2026; Harries, 2023).

RNA processing is one of a collection of phenomena that mediate the lifecycle of an RNA, and the RNA binding proteins (RBP) involved are implicated in multiple aspects of this. Accordingly, they are now recognized as a new traction point in the therapeutic landscape (Neil et al., 2022). The ability to regulate the transcriptome in a labile and responsive manner is a pre-requisite to the plastic and adaptable transcriptome necessary for cellular homeostasis. Alternative splicing (the production of multiple isoforms from a single coding unit depending on the internal and external environment) and correct regulation of RNA turnover are central to this. Many RNA binding proteins involved in splicing decisions also have roles in other processes relevant to senescence such as telomere maintenance, regulation of proinflammatory cytokines and RNA quality control (Latorre and Harries, 2017; Pont et al., 2012). Disruption to RNA processing is associated with cellular senescence and adverse ageing outcomes in vitro and in vivo (Holly et al., 2013; Latorre et al., 2017; Latorre et al., 2018a; Lee et al., 2019; Lye et al., 2019). The expression of RBPs such as splicing factors declines with age because of repeated and constitutive activation of the AKT and ERK signaling pathways, and their effector genes FOXO1 and ETV6 (Latorre et al., 2019). Splicing factor expression can be restored by naturally occurring small molecules such as polyphenols (Latorre et al., 2017) or H_2_S donors (Latorre et al., 2018b), or by inhibition of their upstream negative regulators AKT and ERK (Latorre et al., 2019). These interventions result in reprogramming of senescent cell populations and the attenuation of the SASP, with or without rebuilding of telomeres and resumption of cell cycle, depending on the point of intervention. Some agents have been shown to protect against the development of senescence in human cell models raising the possibility of prophylactic treatment (Latorre et al., 2018b).

Here, we sought to identify and evaluate novel combinations of natural and nature-identical synthetic senomorphic molecules capable of reprogramming senescent skin cells via the AKT/ERK/splicing regulation axis in naturally aged human primary 2D and 3D skin models for development as skin health actives. We demonstrate here the beneficial reprogramming impact of naturally-derived small molecules in targeting skin cell senescence in primary human systems and propose that such approaches comprise a novel and effective approach to skin health, by targeting the upstream effectors, rather than the downstream consequences of skin ageing.

## Results

### AKT and/or ERK inhibitors target splicing regulation

Inhibitors of the ERK and AKT pathways have previously been documented to inhibit the expression of splicing factors and reprogram senescent cells (Latorre et al., 2019). We identified 69 nature-identical compounds with literature or computational evidence of interaction with ERK and/or AKT. We tested a subset of 14 molecules for impact on the expression of a panel of previously implicated splicing regulator genes in senescent primary human dermal fibroblasts by qRT-PCR. Splicing factor expression is autoregulated and labile, but we demonstrated that even at a single timepoint, all 14 molecules had dynamic and robust impact on the expression of at least one splicing factor, and most had impact on multiple splicing factors (figure 1).

**Figure 1:**
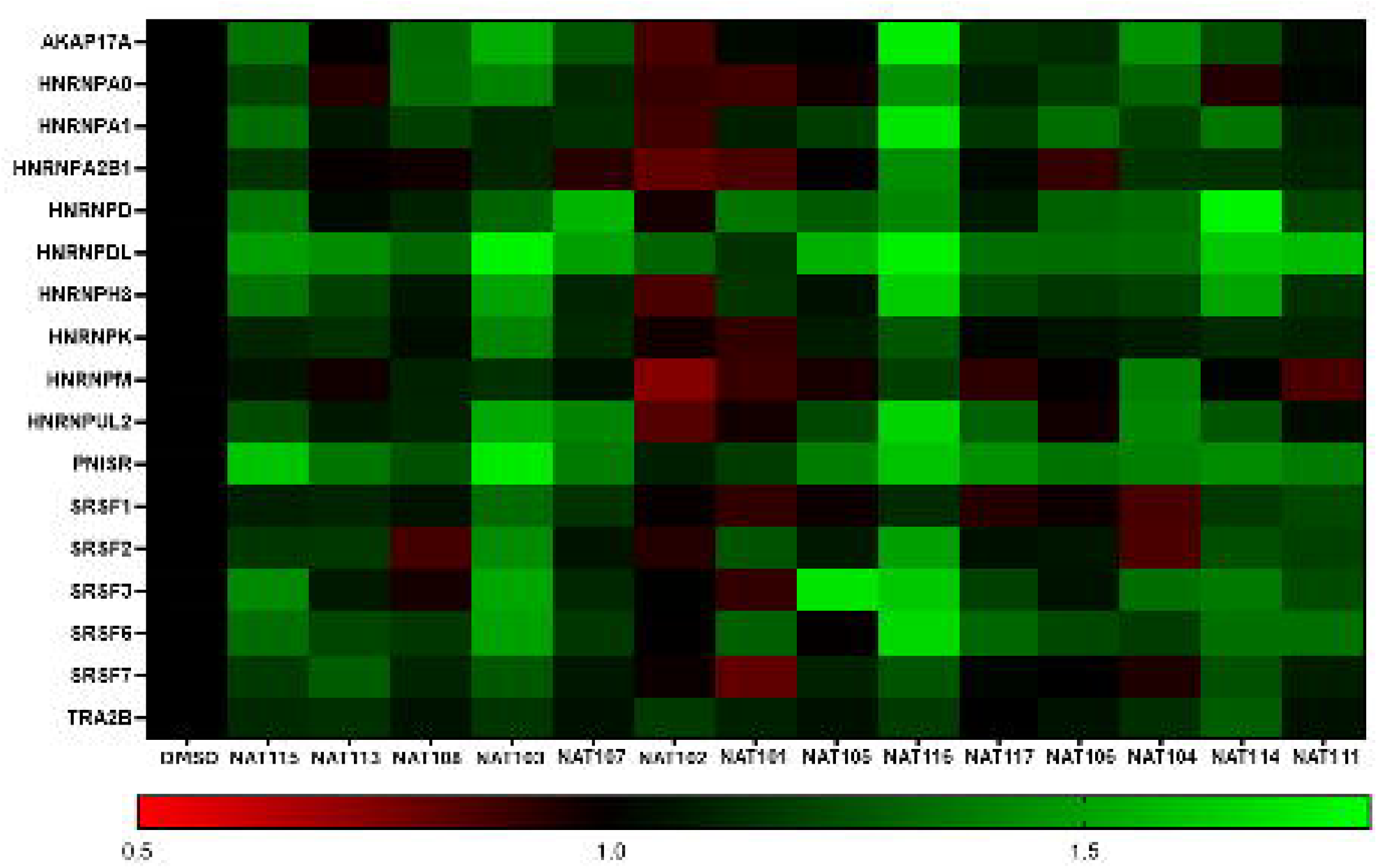
The impact of ERK or AKT inhibitors on the expression of splicing regulators. Small molecules selected by shape/charge similarity, similar molecular docking characteristics to inhibitors or known relationships with target pathways cause changes to the expression of splicing regulator transcripts. Splicing factor identity is given on the y axis and molecule identity on the X axis. Quantifications are fold change relative to the DMSO-only control, derived from qRT-PCR quantification. Green boxes refer to upregulation relative to controls whilst red refers to down-regulation.

### 82 combinations of nature-identical synthetic molecules have synergistic impact on senescent cell load in human primary dermal fibroblasts in vitro

We assessed all possible combinations of 69 nature-identical AKT and/or ERK inhibitors for impact on senescent cell load in aged male primary dermal fibroblasts in vitro. 148 combinations demonstrated senolytic activity at the doses used (mostly involving NAT104, NAT151, NAT155 and NAT168) in cultured fibroblasts. 331 combinations showed evidence of senomorphic activity (reduction in senescent cell load with minimal impact on cell number), of which 82 demonstrated evidence of synergy (effects ≥10% greater than the sum of the individual molecules). The results of treatment with the top 10 best performing combinations assessed by drop in senescent cell load and observed synergy are presented in figure 2.

**Figure 2:**
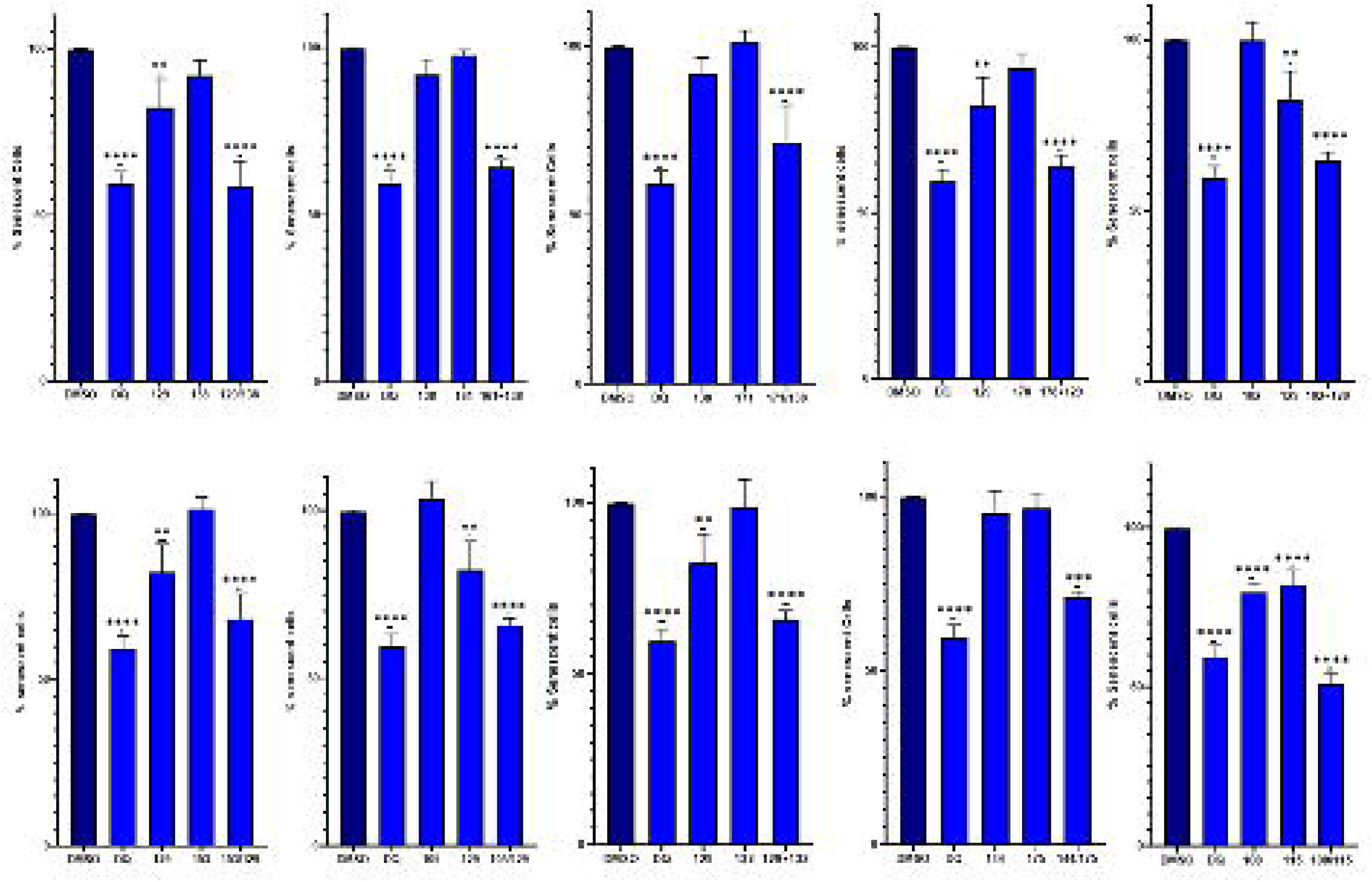
Senomorphic small molecule combinations in naturally aged primary human dermal fibroblasts. The percentage of senescent cells present in a naturally aged primary human dermal fibroblast culture is given. DMSO is the vehicle only control, DQ = dasatinib and quercetin positive control. Data were normalized to the percentage of senescent cells in the DMSO vehicle control. Statistical significance is by one way ANOVA. * = p<0.05, ** = p<0.01, ***=p<0.001, **** = p<0.0001.

### Dose ranging studies and cell kinetic analysis of top 10 compound combinations in senescent male and female primary human fibroblasts

Best performing combinations were validated in cultures of both male and female naturally-aged human primary fibroblasts. All 10 combinations demonstrated evidence of senomorphic effect in primary cells of both sexes. Combination NAT115/NAT108 did not perform as well in high throughput assay on male or female cells and was discounted from further study. Most compounds showed stability of effect over the dose range tested (figure 3). No overt differences in response between male and female cells were noted (figure 4). Cytotoxicity as measured by

**Figure 3:**
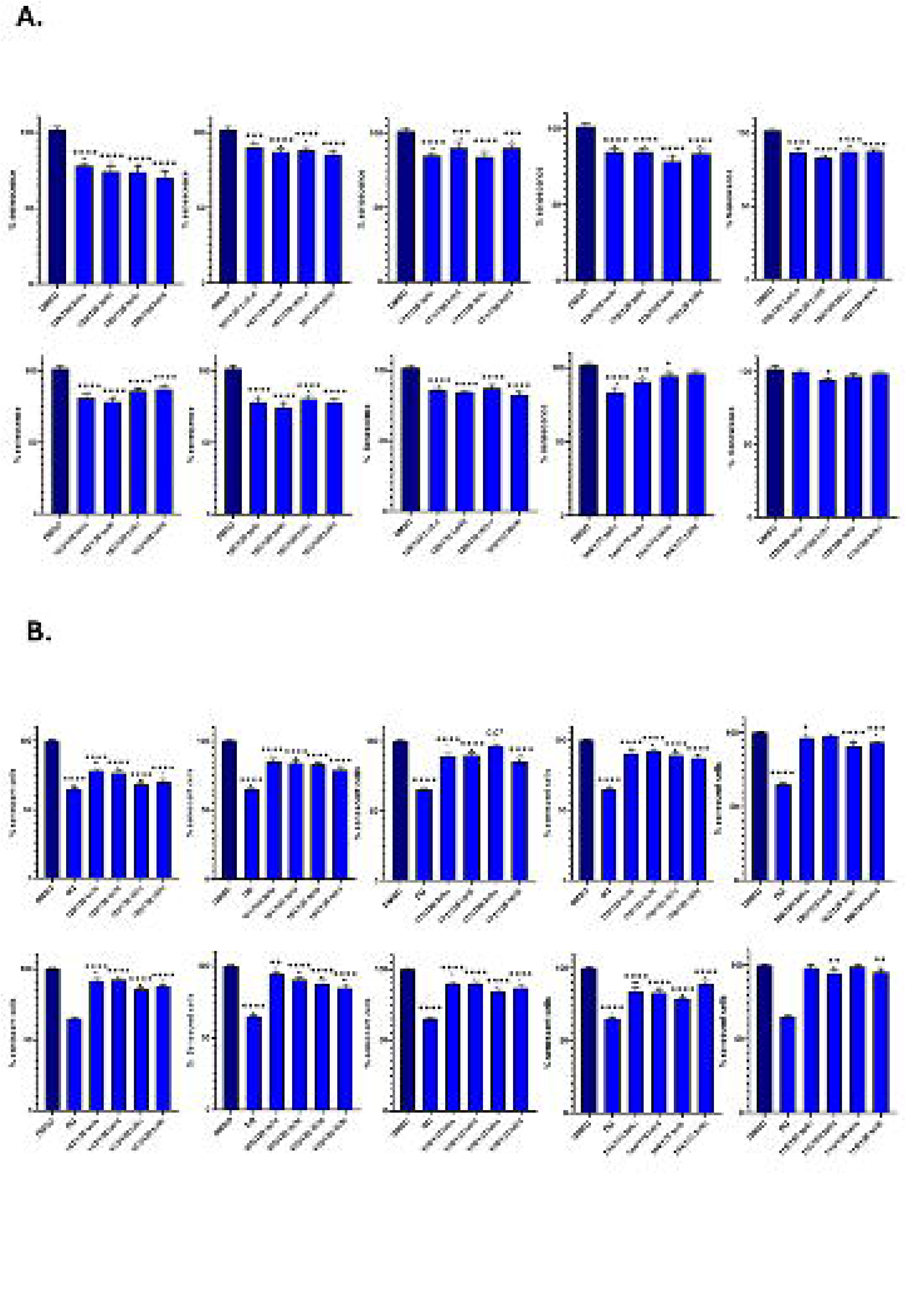
Impact of compound combination dose on senomorphic activity in male (A) and female (B) human primary dermal fibroblasts. Compound combinations were tested over a range of four empirically determined doses. Senescent cell load is normalized to the vehicle-only control and compared to the established senolytic combination dasatinib and quercetin (DQ). Statistical significance is by one way ANOVA. * = p<0.05, ** = p<0.01, ***=p<0.001, **** = p<0.0001.

**Figure 4:**
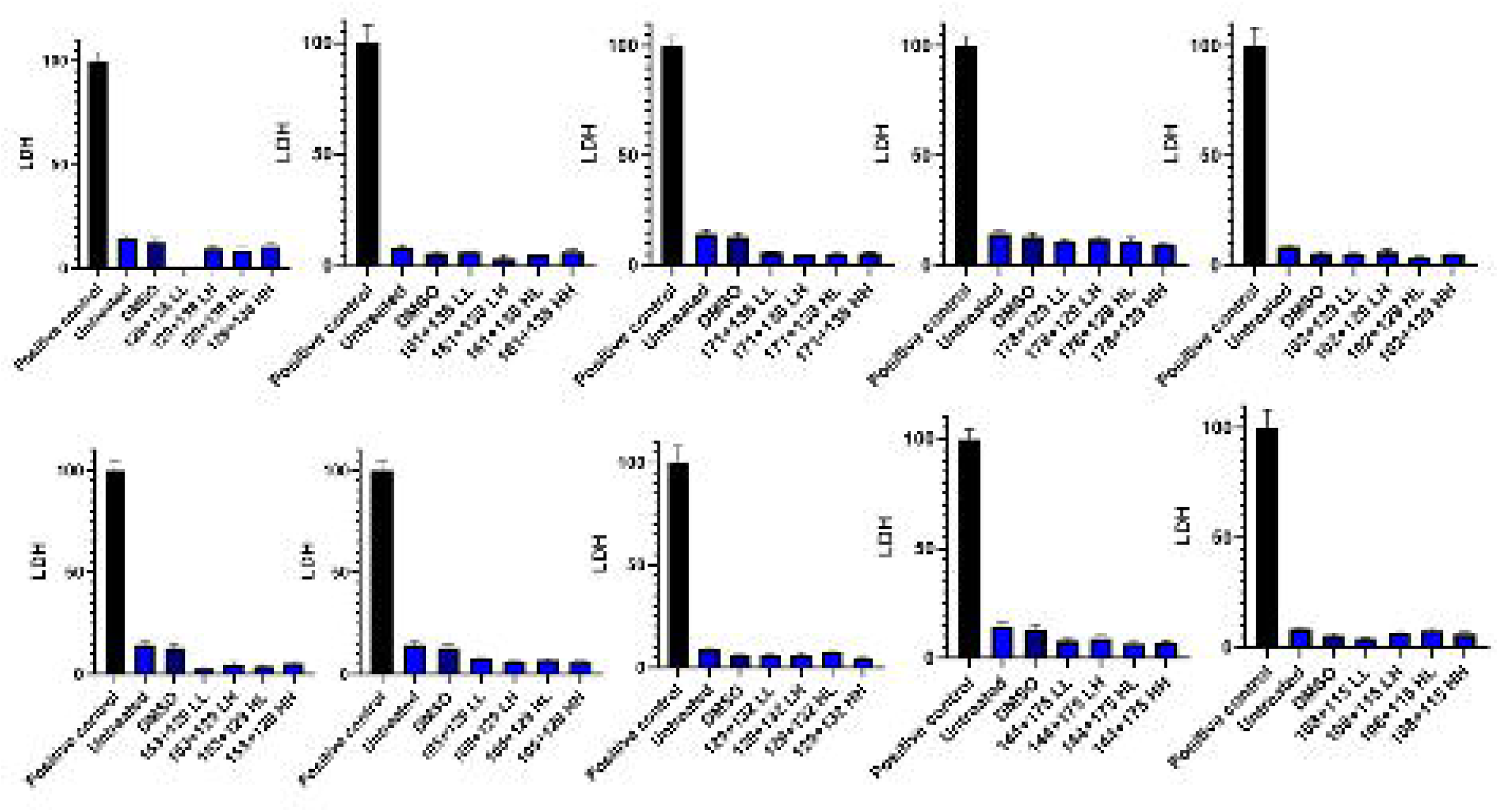
Cytotoxicity of compound combinations assessed in senescent male primary human dermal fibroblasts by LDH assay. The impact of four different doses of compound combinations on toxicity as measured by LDH assay is given relative to the positive control (cell lysis buffer provided in kit) and the DMSO vehicle control. Data are presented normalized to positive control. LL = Low/Low, LH = Low/High, HL = High/Low and HH = High/High.

LDH assay revealed that none of the compound combinations exhibited any significant toxicity in this system at the doses used.

### Natural extracts containing identified actives (or related derivatives) produce similar senomorphic effects

Seven compound combinations were selected for further development, prioritizing those with actives (or derivatives) found in natural plant extracts that could be readily sourced. These extracts were then used for subsequent experiments. Of those, 5 combinations demonstrated an impact on features of senescence including reduction in senescent cell load, reduction in induced proliferation, reduction in DNA damage load and/or reduction in SASP markers (table 1). Combinations A & C and B & C were discounted because of apparent increases in the inflammatory marker IL-8, whereas the combination of G & H was discounted due to lack of impact on senescence in keratinocytes. The combination of extract A and extract E (*H. lupulus* and *G. biloba*) was selected as the final development candidate (figure 5). This combination of extracts produces a significant dose-dependent reduction in senescent cell load of up to 45% and 10% in aged primary human dermal fibroblasts (p = <0.0001) and keratinocytes (p = <0.001) respectively, with no impact on resumption of proliferation in either cell type (figure 5). A 50% drop in DNA damage load was also observed in aged primary human dermal fibroblasts at lower doses (p = <0.0001). Characteristic changes in morphology relating to reprogramming of senescent cell functionality were also evident in both dermal fibroblasts and keratinocytes (figure 5).

**Figure 5:**
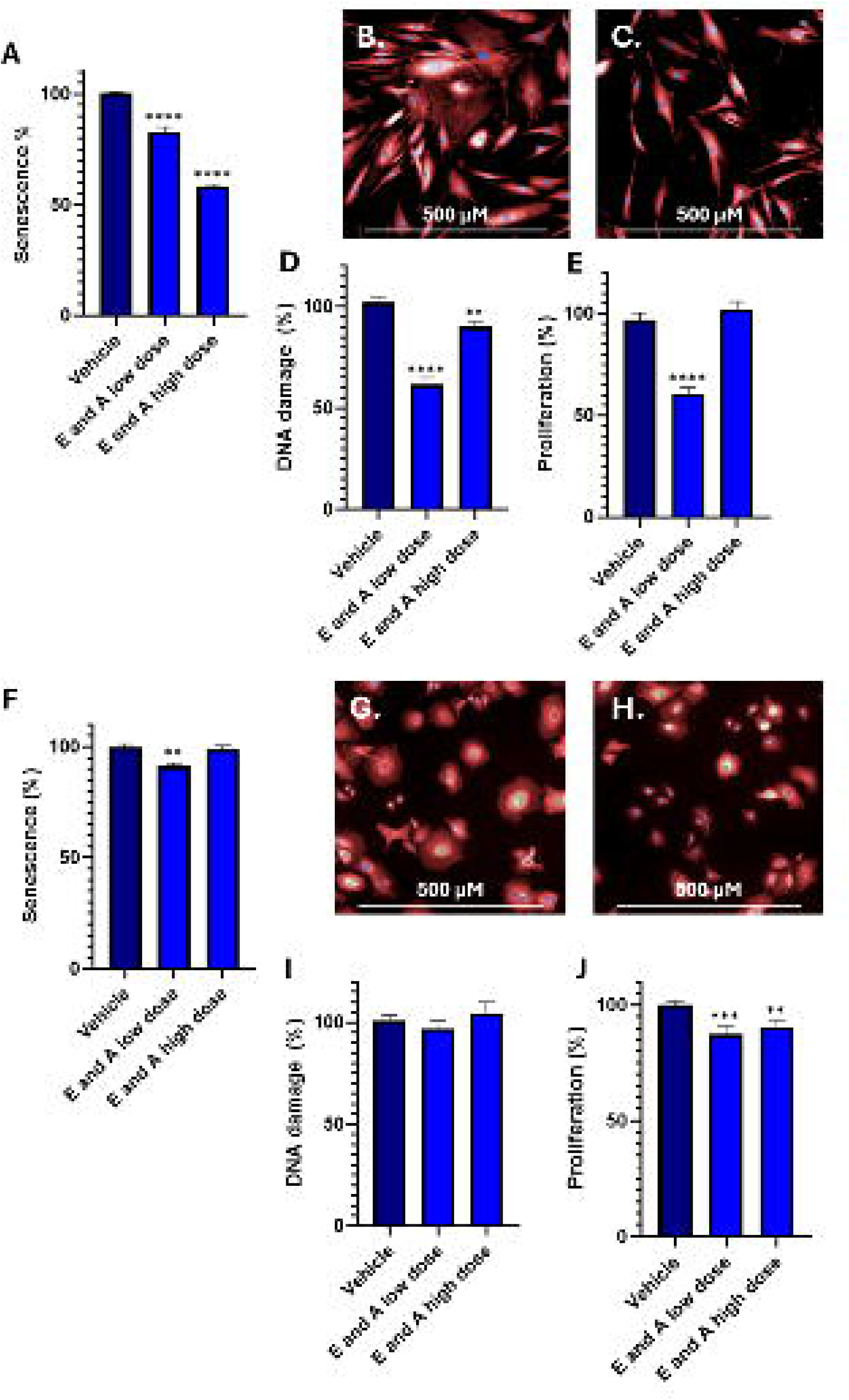
A combination of extracts A and E demonstrates a senomorphic action in senescent primary human dermal fibroblasts and keratinocytes. **A-E** = dermal fibroblasts, **F–J** = keratinocytes. **A** = senescent cell load, **B** = untreated senescent primary human dermal fibroblasts, **C** = treated primary human dermal fibroblasts, **D** = DNA damage load, **E** = Proliferation, **F** = senescent cell load, **G** = untreated senescent primary human keratinocytes, **H** = treated primary human keratinocytes, **I** = DNA damage load and **J** = proliferation. Nuclei are shown in blue, cytoplasm in red and SA-β-Gal in green. Statistical significance is by unpaired two-sided t test. * = p<0.05, ** = p<0.01, ***=p<0.001, **** = p<0.0001.

**Table 1:** Five combinations of natural extracts demonstrate impact on features of senescence in naturally-aged human primary dermal fibroblasts and keratinocytes.

| Extract 1 | Extract 2 | Cell Type | Senescence | Proliferation | DNA damage | SASP |
| --- | --- | --- | --- | --- | --- | --- |
| A | E | Fibroblasts | Reduced | Reduced | Reduced | No Change |
|  |  | Keratinocytes | Reduced | Reduced | Reduced | No Change |
| A | D | Fibroblasts | Reduced | Reduced | Reduced | Reduced |
|  |  | Keratinocytes | Reduced | Reduced | No Change | Reduced |
| A | C | Fibroblasts | Reduced | Reduced | No Change | Increased |
|  |  | Keratinocytes | Reduced | No change | Reduced | Increased |
| B | C | Fibroblasts | Reduced | Reduced | No Change | Reduced |
|  |  | Keratinocytes | Reduced | Reduced | No Change | Increased |
| G | H | Fibroblasts | Reduced | Reduced | Reduced | No Change |
|  |  | Keratinocytes | No change | No Change | No Change | Reduced |

### Natural extracts containing senomorphic actives result in restoration of skin structure in 3D human skin models

Treatment with a simple formula including a combination of *H. lupulus* and *G. biloba* produced qualitative improvement in features of senescence, skin substructure and inflammation. Compared to untreated controls, UV irradiation alone provoked hyperplasia of the epidermis, disorganization of the collagen substructures, and increases in the senescence marker p21 and inflammatory markers IL-6 and IL-8. In contrast, in samples treated with the combination of extracts A and E following UV irradiation, qualitative evaluation of these parameters revealed a reduction in p21 (p = 0.06), a validated biomarker of senescence, resolution of epidermal hyperplasia as evident on H&E staining, normalization of collagen fiber structure in the dermis visualized by changes in Masson’s Trichrome staining, as well as a reduction in IL-8 visualized by immunofluorescence (figures 6 and 7). Although the variability, as expected in human primary skin samples, p21 data did not quite reach statistical significance (p = 0.06), however these data are consistent with qualitative reduction in senescence and consequent improvement in skin structure following topical treatment in simple formulation.

**Figure 6:**
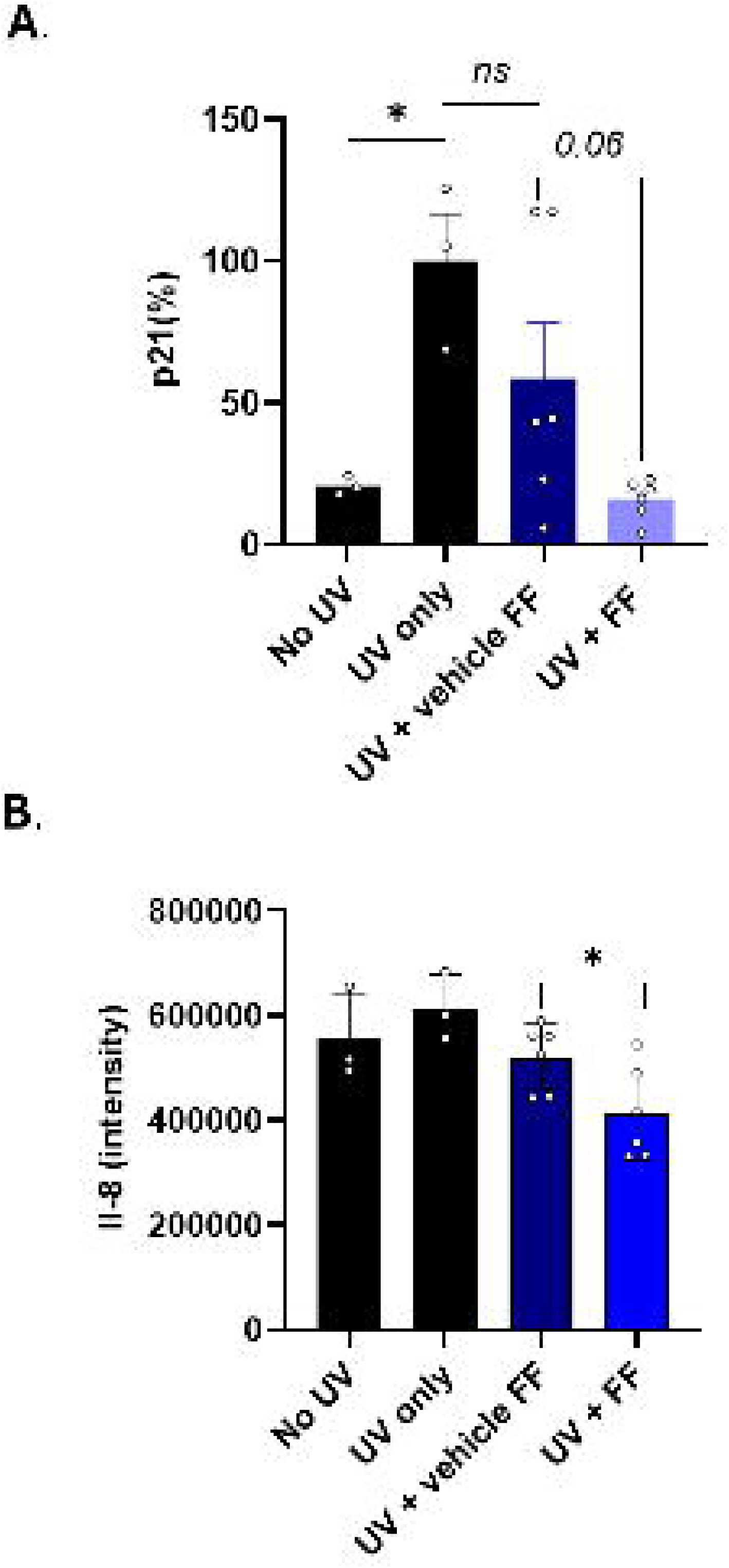
Graphical representation of p21 and Il-8 expression in UV-treated full thickness 3D human skin. Both p21 and IL8 can be seen to trend downward following treatment of UV treated skin samples with topical application of simple formulation. Inter-sample variability in the control treated samples has however resulted in reduced power for statistical significance for p21. FF = simple formulation. UV = UV-treated control.

**Figure 7:**
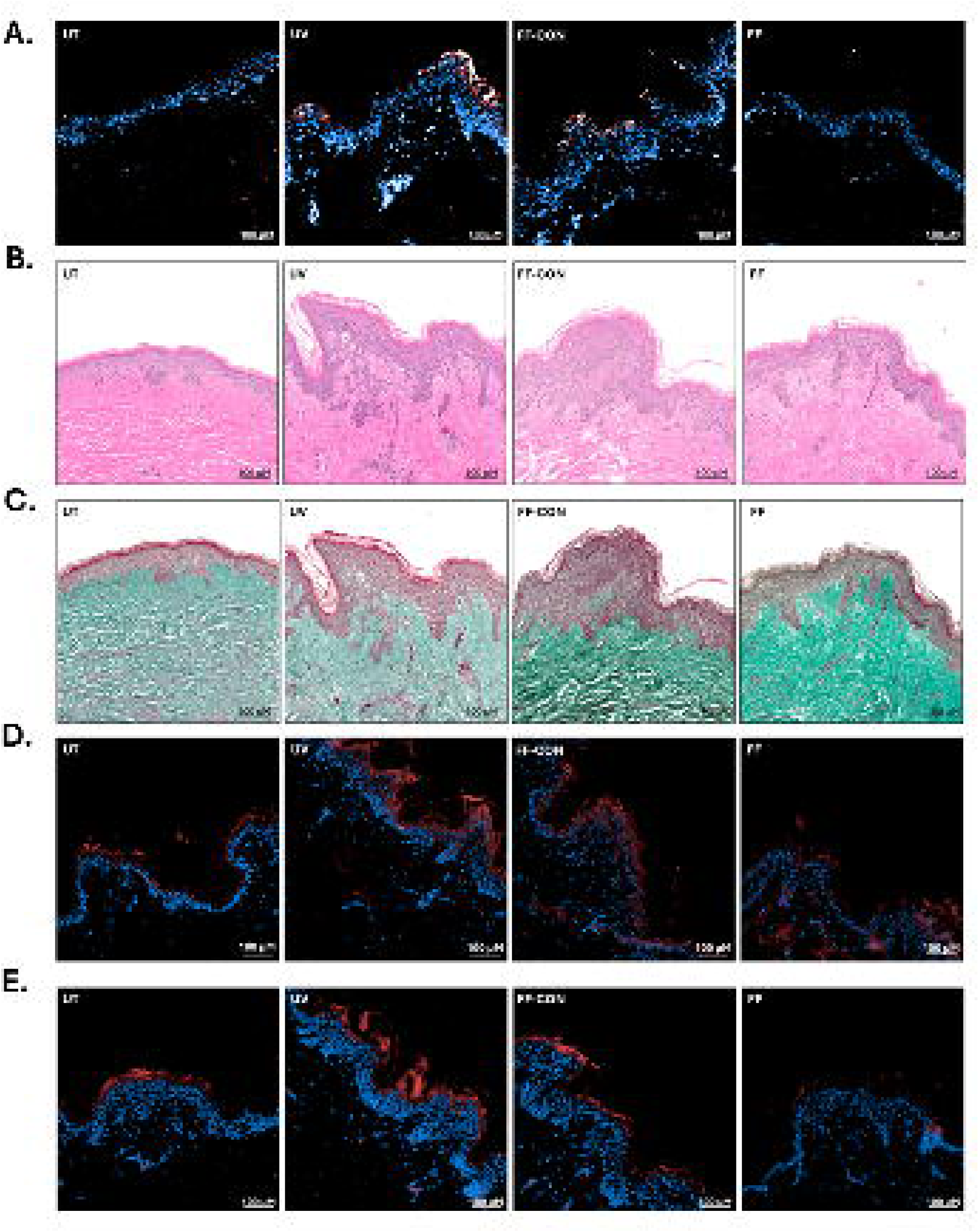
The impact of topical application of a combination of extracts A and E on skin structure, senescence and inflammation. **A** = p21 staining for senescence. A reduction in p21 is consistent with a reduction in senescent cell load. **B**. Haematoxylin and eosin (H&E) staining for skin substructure. **C**. Masson’s Trichrome staining for extracellular matrix proteins. Collagen fibers are picked out in jade. **D.** IL-6 immunofluorescence and **E**. IL-8 immunofluorescence. UT = Untreated control. UV = UV-treated control, FF-CON = control for simple formulation. FF = topically applied simple formulation. Scale bars are 100μm.

## Discussion

We aimed to identify and evaluate naturally derived or nature identical small molecules for development as novel skin health actives. We evaluated combinations of natural and nature-identical synthetic small molecule ERK and/or AKT inhibitors for impact on senescence in naturally aged primary human dermal fibroblasts and keratinocytes, and in complex human ex vivo 3D full thickness skin explants. We identified 331 combinations of molecules with reprogramming activity, 82 of which demonstrated a synergistic mode of action. 10 combinations had similar efficacy in male and female cells across doses with no cytotoxic effects noted in either sex. 5 combinations retained efficacy when actives were delivered in the form of plant extracts. These data enabled prioritization of a combination of *H. lupulus* and *G. biloba* extracts as the development candidate which caused a reproducible decrease in senescent cells and DNA damage burden in treated cell populations, together with the absence of resumption of cell proliferation. We also observed near significant reduction in senescence biomarkers (p = 0.06) and beneficial action on skin substructure and inflammation when delivered to human 3D skin explants topically in simple formulation. These data suggest that a combination of *H. lupulus* and *G. biloba* extracts delivered topically could reprogram senescent human cells and improve the quality and health of skin when utilized in a skincare setting.

There is currently great interest in the use of agents that can remove or reprogram senescent cells in the context of skincare. A reduction of only 25-30% in senescent cell load has been shown to result in clinical benefit (Justice et al., 2019; Pignolo et al., 2020). The magnitude of impact we have demonstrated here lies well within range upon a single application and will likely be larger with repeated administration (Latorre et al., 2017). Removal of senescent cells by senolysis, particularly with first generation senolytics, may eliminate multiple senescent cell subtypes without discrimination, including those which may be beneficial (Attaallah et al., 2019). Large scale apoptosis may also cause tissue damage and contribute to disease pathogenesis (Rock and Kono, 2008). In contrast, a senomorphic approach retains cells in their proper context and should preserve tissue-resident stem cells for improved tissue rebuilding.

Targeting senescence via the splicing regulatory axis provides a unique ability to target this process within the normal homeostatic parameters of the cell and without the resumption of senescent cell proliferation. Splicing factors are labile and responsive molecules that respond to the internal and external environment and prime the cell to retain or regain homeostasis even in the face of challenge. Restoration of these proteins has been demonstrated to restore splicing homeostasis and functionality to senescent cells of different lineages (Latorre et al., 2017; Latorre et al., 2019; Latorre et al., 2018b). We have identified a combination of molecules, and a respective combination of extracts (*H. lupulus* and *G. Biloba*) with well-defined safety and tolerability parameters, which can attenuate this axis and bring about a robust decrease in senescent cells and DNA damage load in naturally aged human primary skin cells in vitro. Qualitative changes in skin morphology and structure in 3D primary full thickness human skin explants were also suggested by observations. The use of known extracts brings several advantages. The molecules involved have known and validated human clinical and ecological safety, as well as favorable tolerability parameters; indeed, many of them have been in previous use in other contexts and have proven safety in topical or oral formulations. In the context of skincare product development, regulatory approvals, suppliers and infrastructure around their usage in a cosmeceutical context are well established. Delivering the actives in combination brings an additional and unexpected synergy of effect to maximize utility. There is huge current interest in developing ‘geroscience’ approaches to improve the health of ageing cells. Harnessing the new biology we have discovered represents a novel approach to skincare, which restores the ability of the skin and its substructure to respond to the internal and external challenges which provoke skin ageing.

## Methods

### Cell lines and culture conditions used for study

#### Fibroblasts

Primary human male dermal fibroblasts (nHDF) from the abdomen of a 36yr old Caucasian donor (Cat. #C-12302, Lot. #445Z026.3) and primary human female dermal fibroblasts from the breast of a Caucasian 28yr old donor (Cat. #C-12302, Lot. #467Z026.3) were purchased from PromoCell GmbH (Heidelberg, Germany). Cells were seeded in T75 flasks at a density of 3 X 10^5^ cells per culture and brought to senescence by continuous culture until they reached a population doubling time of at least 136 hours and displayed the characteristic increase in cell size indicative of a senescent cell population. Cells were maintained in Fibroblast Growth Medium 2 (Cat. # C-23220; PromoCell GmbH, Heidelberg, Germany) supplemented with Fibroblast Growth Medium 2 Supplement Mix (Cat. # C-39325; PromoCell GmbH, Heidelberg, Germany), and 1% Penicillin/Streptomycin, incubated at a temperature of 37°C with an atmospheric CO2 level of 5%, and passaged whenever cell confluency reached 90%. Penicillin-streptomycin was removed during the final passage immediately preceding experimental treatment.

#### Keratinocytes

Two primary human female keratinocyte cell lines were purchased from Promocell GmbH, Heidelberg. Line 1 was from the abdomen of a 56yr old donor (Cat. #C-12003, Lot. #451Z014.3) whereas line 2, used for validation only, was from the temple of a 58yr old Caucasian woman (Cat. #C-12003, Lot. #489Z018.2). Cells were seeded in T75 flasks at 4 X 10^5^ cells per culture and maintained at 37°C and 5% CO_2,_ in serum-free Keratinocyte Growth Medium 2 containing Keratinocyte Basal Medium 2 (PromoCell, Cat. #C-20211), Keratinocyte Growth Medium 2 Supplement Pack (PromoCell, Cat. #C-39011) and 0.06 mM CaCl_2_. Penicillin-streptomycin (Gibco, Cat. #15140122; Thermo Fisher Scientific, Waltham MA, USA) was included in the medium for all routine culture phases but was removed during the final passage immediately preceding experimental treatment.

### Selection of molecules for study

Nature-identical molecules were selected for screening by documented interaction or shape/charge similarity and docking characteristics comparable with known specific AKT or ERK inhibitors. Computational chemistry was undertaken by Domainex (www.Domainex.co.uk), whilst other molecules were included because of evidence in the literature. Molecules were coded NAT101 – NAT165. Molecules were purchased from A2B Chem LLC, AA Blocks LLC, AnalytiCon Discovery GmbH, BLD Pharmatech GmbH, MedChemExpress Europe AB, Merck KGaA, Molport SIA, TargetMol Chemicals Inc, and TimTec LLC.

### Assessment of impact on splicing factor expression

A selection of 14 molecules was evaluated for impact on splicing factor expression by high-throughput RT-qPCR using custom-designed OpenArray® plates on the Thermo Fisher QuantStudio 12K Flex platform (Thermo Fisher Scientific, Waltham MA, USA). Reaction mixes contained 2.5 μl 2 OpenArray® Real-Time Master mix (Thermo Fisher Scientific, Waltham MA, USA), cDNA (1.2μL) and RNase-free dH_2_O (1.3μL) (Thermo Fisher Scientific, Waltham MA, USA). Gene expression was measured by relative quantification normalized to the geometric mean of three housekeeping genes (*IDH3B*; Hs00199382_m1], *POLR2A; Hs00172187_m1]* and *TMEM199*); Hs01022209_m1) and expressed relative to vehicle only control. TaqMan assay Ids (Thermo Fisher Scientific, Waltham MA, USA) are indicated in parentheses. Experiments were carried out in 3 biological replicates and 3 technical replicates. Differential expression in treated and untreated cells was then assessed by one-way ANOVA using GraphPad Prism V10 (GraphPad Software, Boston, USA).

### High throughput screen for senomorphic activity

69 nature-identical synthetic molecules were identified for evaluation in a primary screen of 2346 pairwise combinations for impact on senescence phenotypes in male human primary dermal fibroblasts. Cells were seeded at a density of 750 cells per well in Revvity 384-well PhenoPlates (Revvity, Cat. # 6055302, Waltham MA, USA). Each test substance was reconstituted in DMSO for a stock concentration of 100mM. The stock solutions were prepared in DMSO and used at a final concentration of 2.5μM. Combinations and individual molecules were tested in triplicates at 2.5μM. Compounds were designated as senomorphic rather than senolytic, if reprogramming occurred in the absence of overt cell loss (defined as a >20% decrease in cell number). The known senolytic compounds dasatinib and quercetin (DQ), and DMSO vehicle alone were included as controls. Cells were incubated at 37°C for 24hrs prior to evaluation.

For high content imaging, cells were fixed with 2% formaldehyde + 0.2% glutaraldehyde and washed with PBS prior to staining. Nuclei were stained using 0.5 μg/ml Hoechst working solution (Invitrogen™ Hoechst 33342, Trihydrochloride, Trihydrate - 10 mg/mL Solution in Water; Thermo Fisher Scientific, Waltham MA, USA) for 10 minutes. Senescence-associated beta galactosidase (SA-β-Gal) staining was carried out using the CellEvent™ Senescence Green Detection Kit (Thermo Fisher Scientific, Nuclei were stained using 0.5 μg/ml Hoechst working solution (Invitrogen™ Hoechst 33342, Trihydrochloride, Trihydrate - 10 mg/mL Solution in Water; Thermo Fisher Scientific, Waltham MA, USA) for 10 minutes, Waltham MA, USA) according to manufacturer’s instructions. After incubation cells were washed with PBS prior to the application of CellMask Deep Red staining solution (Invitrogen™ CellMask™ Deep Red Plasma Membrane Stain; Cat. # H32721, Thermo Fisher Scientific, Waltham MA, USA) with incubation for 30 minutes at 37°C with no CO_2_. Following staining, cells were imaged using an Operetta CLS™ High Content Analysis system (Revvity, Waltham MA, USA), and images analyzed using the Harmony™ PhenoLOGIC™ High-Content Imaging and Analysis Software version 5.1 (Revvity, Waltham MA, USA). Senescent cell load was determined using a proprietary multiparameter machine-learning-informed algorithm that integrates cellular parameters such as cell size, cell morphology, nuclear size, nuclear morphology and SA-β-Gal staining to categorize and quantify senescent cell load. The algorithm was validated in-house and externally (www.selvita.com). 10 combinations of molecules were prioritized for secondary screening according to an effect size of >25% for senescence, evidence of ≥10% synergy, minimal decrease in cell number and favorable physicochemical characteristics for future formulation.

### Characterisation of senomorphic effect across dose in naturally aged male and female primary human dermal fibroblasts

Prioritized combinations of compounds were characterized across multiple doses in both male and female primary human cells. Senescent human dermal fibroblasts were seeded onto Revvity optically-clear 96-well PhenoPlates (Revvity, Cat. # 6055302, Waltham MA, USA) at 2000 cells/well and treated with prioritized combinations of nature-identical synthetic small molecules over four dose combinations; low dose of both molecules, low dose of molecule 1/high dose of molecule 2, high dose of molecule 1/low dose of molecule 2, high dose of both molecules (supplementary table 1) was carried out in 8 replicates for 24 hours. Imaging and analysis were conducted as described above. Cytotoxicity was assessed by measuring the release of cytoplasmic lactate dehydrogenase (LDH) into the culture supernatant using the CytoTox 96® Non-Radioactive Cytotoxicity Assay (Promega, Cat. #G1780 Madison WI, USA) according to manufacturer’s instructions.

### Evaluation of natural extracts containing SENISCA’s actives in 2D primary cultures of senescent primary human dermal fibroblasts

We identified 36 natural plant extracts containing the same molecules to those used in our synthetic combinations, or close derivatives thereof. From these, 7 combinations were selected based on our earlier in vitro data but also considering existing safety and toxicology data, compliance with existing regulatory protocols and known relationships with validated vendors (supplementary table 2). Dosing for each replacement extract was established empirically by lack of observable cytotoxicity and similar impact to previously tested synthetics on senescent cell load. Senescent female primary human dermal fibroblasts and keratinocytes were seeded at 2000 and 4000 cells per well, respectively, in Revvity optically-clear 96-well PhenoPlates (Revvity, Cat. # 6055302, Waltham MA, USA) and treated with each combination of test extracts or related controls and incubated for 24 hours prior to analysis. Impact on senescent cell load and cytotoxicity were assessed as described above. Cell proliferation kinetics and DNA damage were measured in multiplex; after fixation with 2% formaldehyde + 0.2% glutaraldehyde, cells were fluorescently labelled using Hoescht 33342 (Thermo Fisher Scientific, Cat. # 62249, Waltham MA, USA) to define the cell nuclei, Alexa Fluor® 488 Anti-Ki67 (Abcam, Cat # ab558616; Cambridge, UK) at 1:750 dilution for proliferation and Alexa Fluor® 647 Anti-gamma H2A.X (phospho S139) antibody (Abcam, Cat. # ab81299; Cambridge, UK) at 1:750 dilution to define DNA-damaged cells. Cells were incubated at room temperature for 1h protected from light.

For SASP immunocytochemical analysis, NHDF and NHEK cells cultured in Revvity optically-clear 96-well PhenoPlates (Revvity, Waltham MA, USA). Fixation and blocking were as previously described. Cells were then incubated for 1 hour at room temperature with primary antibodies against IL-6 (Rabbit anti-IL-6, Abcam, Cambridge, UK, Cat. #ab233706) at 1:50 dilution or IL-8 (Rabbit anti-IL-8, Abcam,

Cambridge, UK, Cat. #ab289967) at 1:100 dilution. Bound primary antibodies were visualized by incubation with an Alexa Fluor® 647-conjugated donkey anti-rabbit IgG secondary antibody (Abcam, Cambridge, UK, Cat. #ab150075) at 1:1000 dilution for 30 minutes at room temperature, followed by a Hoechst counterstain to visualize cell nuclei. Cells were then imaged using an Operetta CLS™ High Content Analysis system (Revvity, Waltham MA, USA), and images analysed using the Harmony™ PhenoLOGIC™ High-Content Imaging and Analysis Software version 5.1 (Revvity, Waltham MA, USA).

### Human ex vivo full thickness skin 3D assessment of action on skin ageing characteristics

The ability of the lead combination to bring about changes in biochemical and histological features of skin ageing was assessed in full thickness 3D human skin models. Skin was procured from GenoSkin (Toulouse, FR) following donor surgery and was maintained for 24hrs for acclimatization prior to experimentation. Samples were supplied with informed consent at source. The donor was a 29yr old female of phototype type 3. Skin samples were cultured at 37°C, 5% CO_2_ and 95% humidity, in proprietary media supplied with the samples. Senescence was induced using total UVC dose of 1.5 J/cm2 using a UV crosslinker (HL-2000 HybriLinker, UVP) to produce experimentally tractable levels of senescence. Actives in simple formulation containing additional proprietary additives were then topically applied to the sample surface and cultured for 24h. The conditioned medium was harvested and stored at - 80°C, and each skin sample was cut into 2 halves for OCT and FFPE embedding steps. Slides were stained with haematoxylin and eosin (H&E) for tissue structure, and Masson’s Trichrome (MT) for collagen and extracellular matrix structures, according to established protocols. Digital image analysis was performed using QuPath (version 0.5.1). Custom colour deconvolution vectors were generated for H&E and MT staining from representative images by manually selecting small regions of interest (ROIs) containing isolated staining signals (e.g., haematoxylin-positive nuclei). The resulting vectors were verified across the image dataset prior to analysis. Quantification was performed using the QuPath threshold-based pixel classification tools with Gaussian prefiltering. Positive pixels were identified using optical density thresholds selected to distinguish target staining from background tissue. Manual annotations were used to define regions for analysis and exclude artefacts and non-tissue areas. Deconvolution and classification settings were subsequently applied to all images using automated batch processing.

For immunohistochemical analysis, air-dried cryosections were fixed in 1% formaldehyde (FA) with 0.1% glutaraldehyde (GA) in PBS for 10 minutes at room temperature (RT). Fixed samples were quenched with 0.1 M glycine in PBS (10 min, RT), washed twice in PBS, permeabilized with 0.3% Triton X-100 (10 minutes) and blocked with 5% normal goat serum in PBS with 0.3% Triton X-100 (10 minutes, RT). Sections were incubated overnight at 4 °C with primary antibodies against p21 [EPR362] (Abcam, Cambridge, UK, Cat. #ab109520) at 1:150 dilution, IL-6 (Abcam, Cambridge, UK, Cat. #ab6672) at 1:250 dilution or IL-8 (Cell Signaling Technology Inc., Danvers MA, USA, Cat. #94407) at 1:100 dilution in 5% normal goat serum with 0.3% Triton X-100. Slides were then washed three times in PBS containing 0.3% Triton X-100 and incubated with Alexa Fluor® 647-conjugated donkey anti-rabbit IgG secondary antibody (Abcam, Cambridge, UK, Cat. #ab150075) at 1:1000 dilution in 5% normal goat serum in PBS with 0.3% Triton X-100, for 1h at room temperature. Slides were counterstained with DAPI and mounted using an anti-fade mounting medium (Invitrogen™ Fluoromount-G™ Mounting Medium, Cat. #00-4958-02; Thermo Fisher Scientific, Waltham MA, USA) and imaged using the Operetta CLS™ High Content Analysis system (Revvity, Waltham MA, USA). Images were analysed using the Harmony™ PhenoLOGIC™ High-Content Imaging and Analysis Software version 5.1(Revvity, Waltham MA, USA).

## Ethics statement

All primary cells and tissues used in this study were commercially procured with ethical agreement at source.

## Data Availability statement

The authors affirm that all data necessary for confirming the conclusions of the article are present within the article, figures, and tables.

## Acknowledgements

The authors are grateful for the collaboration of Jean-Thierry Simmonnet, who provided thoughtful discussion on study design and lead molecule choice. The research was carried out at the National Institute for Health and Care Research (NIHR) Exeter Biomedical Research Centre (BRC). The authors gratefully acknowledge the staff of the NHS Pathology laboratories at the Royal Devon University Hospital trust (NDUH) for aid in processing 3D skin cryosamples.

## Author contributions

**Conceptualization**: Jouy, C, Ballihaut, C, Lee, BP, Harries, LW;

**Data curation**: Perciato, ML, Davies, M, Akhbari, P

**Formal analysis**: Perciato, ML, Davies, M, Akhbari, P

**Funding acquisition**: Cole, SL, Lee, BP, Harries, LW;

**Investigation**: Perciato, ML, Davies, M, Akhbari, P, Denise, H;

**Methodology**: Perciato, ML, Davies, M, Akhbari, P, Denise, H; Lee, BP;

**Project administration**: Clauss, A, Cole, SL, Jouy, C, Ballihaut, C, Hass, KM, Lee, B, Lee, BP, Harries, LW;

**Resources**: Jouy, C, Ballihaut, C, Hass, KM, Lee, B, Lee, BP

**Software**: Perciato, ML, Davies, M, Akhbari, P, Denise, H

**Supervision**: Akhbari, P, Clauss, A, Lee, BP, Harries, LW

**Validation**: Perciato, ML, Davies, M, Akhbari, P

**Visualization**: Perciato, ML, Davies, M, Akhbari, P, Denise, H

**Writing** — original draft: Harries, LW;

**Writing** — review & editing: Perciato, ML, Davies, M, Akhbari, P, Denise, H, Clauss, A, Cole, SL, Jouy, C, Ballihaut, C, Hass, KM, Lee, B, Lee, BP.

## Declaration of Generative AI use

The development of the classification algorithm had an iterative machine-learning component. The authors did not use AI/LLM in any other part of the research process and/or manuscript preparation.

---

**Supplementary table 1:** Prioritised combinations of senomorphic small molecules for development. Optimised doses of individual molecules are given.

| Molecule 1 | Molecule 2 | Molecule 1<br>Low dose | Molecule 1<br>High dose | Molecule 2<br>Low dose | Molecule 2<br>High dose |
| --- | --- | --- | --- | --- | --- |
| NAT129 | NAT138 | 1.25µM | 2.5µM | 1.25µM | 2.5µM |
| NAT161 | NAT138 | 1.25µM | 2.5µM | 2.5µM | 5µM |
| NAT171 | NAT138 | 2.5µM | 5µM | 2.5µM | 5µM |
| NAT178 | NAT129 | 2.5µM | 5µM | 1.25µM | 2.5µM |
| NAT102 | NAT129 | 2.5µM | 5µM | 1.25µM | 2.5µM |
| NAT153 | NAT129 | 2.5µM | 5µM | 1.25µM | 2.5µM |
| NAT105 | NAT129 | 2.5µM | 5µM | 1.25µM | 2.5µM |
| NAT129 | NAT132 | 1.25µM | 2.5µM | 2.5µM | 5µM |
| NAT144 | NAT175 | 2.5µM | 5µM | 2.5µM | 5µM |
| NAT108 | NAT115 | 1.25µM | 2.5µM | 2.5µM | 5µM |

**Supplementary table 2:** Natural extracts containing molecules related to actives identified in the primary screen.

| Synthetic<br>Molecule A | Extract A | Synthetic<br>molecule B | Extract B |
| --- | --- | --- | --- |
| NAT129 | A | NAT138 | B |
| NAT171 | C | NAT138 | B |
| NAT178 | D | NAT129 | A |
| NAT105 | E | NAT129 | A |
| NAT129 | A | NAT132 | F |
| NAT171 | C | NAT141 | G |
| NAT135 | H | NAT141 | G |

