## Supplementary figures and images for "Targeting skin ageing at the source: a novel cellular reprogramming approach using natural extract combinations"

### Supplementary figure 1

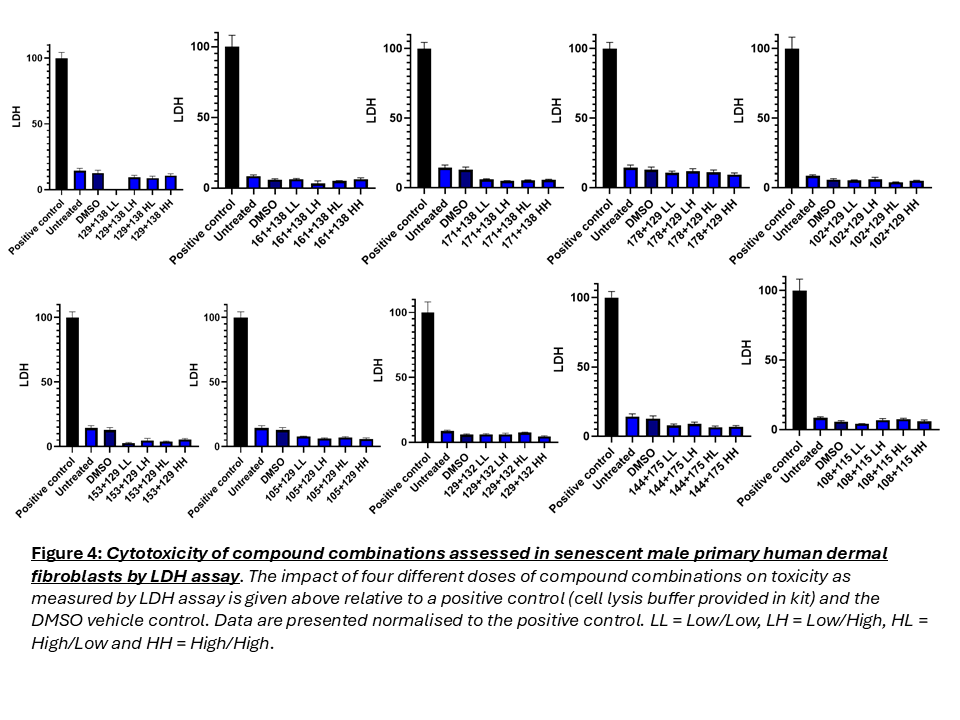
